# Distinct epigenetic shift in a subset of Glioma CpG island methylator phenotype (G-CIMP) during tumor recurrence

**DOI:** 10.1101/156646

**Authors:** Camila Ferreira de Souza, Thais S. Sabedot, Tathiane M. Malta, Lindsay Stetson, Olena Morozova, Artem Sokolov, Peter W. Laird, Maciej Wiznerowicz, Antonio Iavarone, James Snyder, Ana deCarvalho, Zachary Sanborn, Kerrie L. McDonald, William A. Friedman, Daniela Tirapelli, Laila Poisson, Tom Mikkelsen, Carlos G. Carlotti, Steven Kalkanis, Jean Zenklusen, Sofie R. Salama, Jill S. Barnholtz-Sloan, Houtan Noushmehr

**Author notes:** Co-senior author (Jill S. Barnholtz-Sloan, Houtan Noushmehr). CORRESPONDING AUTHOR INFORMATION Houtan Noushmehr, PhD Associate Scientist/Professor Department of Neurosurgery Henry Ford Hospital.

## Abstract

Histomorphology and current grading schemes are unable to predict glioma relapse and malignant tumor progression. We reported that the IDH-mutant associated Glioma-CpG Island Methylator Phenotype (G-CIMP) can be further divided into two clinically distinct subtypes independent of histopathological grading (G-CIMP-high and -low) with evidence of correlation with tumor progression. Here we performed a comprehensive epigenomic analysis of 74 longitudinally collected glioma samples (grade II-IV) to understand malignant recurrence from G-CIMP-high to G-CIMP-low. G-CIMP-low recurrence appeared in 12% of all gliomas and resemble IDH-wildtype primary glioblastoma. G-CIMP-low recurrence can be characterized by distinct epigenetic changes at candidate functional tissue enhancers with AP-1/SOX binding elements, stem cell-like epigenomic phenotype, and genomic instability. Finally, we defined a set of candidate biomarker signatures that predict recurrence of G-CIMP-low with clinically relevance on patient outcomes. Our study provides opportunity for refined clinical trial designs and therapeutic targets that limit progression to more aggressive G-CIMP-low phenotype.

**HIGHLIGHTS:**

- Indolent G-CIMP-high progresses to aggressive G-CIMP-low phenotype
- Incidence of G-CIMP-low recurrent tumors are 3 times greater than G-CIMP-low primary
- G-CIMP-low recurrent tumors share epigenomic features with IDH-wildtype primary GBM
- Predictive biomarkers of G-CIMP-low progression at primary diagnosis

## INTRODUCTION

Heterozygous gain-of-function mutations in *IDH1/2* (Isocitrate dehydrogenase (NADP(+) 1/2) (IDH) is a hallmark of a subset of gliomas associated with favorable patient outcome (Parsons et al., 2008; Yan et al., 2009). Mutant IDH protein produces the oncometabolite 2-hydroxyglutarate (2HG) which may establish the Glioma-CpG Island Methylator Phenotype (G-CIMP) (Noushmehr et al., 2010) by presumably extensive remodeling of the tumor methylome (Turcan et al., 2012). The incorporation of IDH mutation status to the traditional histology by the updated 2016 World Health Organization (WHO) classification of tumors of the Central Nervous System (CNS) represents an emerging concept in which diagnoses of gliomas should be structured and refined in the molecular taxonomy era (Louis et al., 2016).

Glioblastoma (GBM) is a highly aggressive brain cancer and accounts for 46.6% of primary malignant brain tumors with five year overall survival estimate post-diagnosis of 5.5% (Ostrom et al., 2016). Treating recurrent lower-grade glioma (LGG) that undergo malignant tumor progression to GBM is one of the greatest challenges in neuro-oncology (Stupp et al., 2005; Stupp et al., 2009). Currently, the local regrowth of LGG after surgery and adjuvant radio- and chemo-therapies (tumor recurrence) and their malignant progression to GBM is highly variable and unpredictable by histomorphology and grading scheme (Louis et al., 2016). For glioma therapy to be successful, a comprehensive catalogue of molecular biomarkers that can identify a phenotype-shift in gliomas driving tumor recurrence or progression is needed.

Evaluation of genomic heterogeneity across multisector and longitudinal gliomas is informative for targeted therapeutic decisions (Lee et al., 2017). Evidence is emerging that epigenetic abnormalities recapitulate somatic mutation events on cell cycle networks throughout relapse and malignant progression of LGG IDH-mutant G-CIMP glioma cells to GBM (Mazor et al., 2015). Thereby, we sought to define candidate epigenetic features of glioma recurrence deregulated earlier in the development of IDH-mutant non-codel cells harboring the G-CIMP phenotype.

Epigenetically-based molecular classification of 932 primary adult diffuse gliomas (WHO grades II-IV) analyzed by our group have highlighted the existence of three cohesive molecular subgroups of IDH-mutant gliomas (Codel, G-CIMP-high, and G-CIMP-low) and four subgroups of IDH-wildtype (Classic-like, Mesenchymal-like, LGm6-GBM, and PA-like) gliomas with distinct patient outcomes. Accordingly, IDH-mutant DNA methylation signatures allowed the segregation of G-CIMP tumors into two discrete disease subsets independent of neuropathological grading. G-CIMP-low subgroup accounts for 6% of IDH-mutant primary gliomas and is characterized by DNA demethylation and unfavorable survival when compared with the G-CIMP-high subgroup, which accounts for 55% of IDH-mutant primary gliomas (Ceccarelli et al., 2016). IDH mutation is retained throughout glioma recurrence (Mazor et al., 2015; Bai et al., 2016) and using a small cohort of matched primary and recurrent gliomas we have recently reported that a subset of newly diagnosed G-CIMP-high progresses to a G-CIMP-low phenotype (Ceccarelli et al., 2016). However, the critical question of whether G-CIMP-high to G-CIMP-low progression can be determined from a larger cohort and if we can predict G-CIMP-low progression from primary G-CIMP-high gliomas remains unresolved. Here we pooled the largest collection of available genome-wide DNA methylation data from our own study as well as recently published (Mazor et al., 2015; Bai et al., 2016) DNA methylation cohort of longitudinally IDH-mutant and IDH-wildtype glioma samples derived from 74 patients (n=181 adult diffuse glioma fragments WHO grades II-IV) (Tables 1, 2 and S1) as well as available ancillary RNA sequencing, whole-genome sequencing and whole-exome sequencing datasets to further understand glioma recurrence evolution. The IDH-wildtype glioma cases did not change dramatically in terms of their epigenomic profile, but among the IDH-mutants, we defined extensive epigenetic shifts among 12% of all primary G-CIMP-high progressing to the G-CIMP-low phenotype at recurrence, validating and expanding our previous observation. We did not identify any difference between primary *(de novo)* G-CIMP-low and recurrent (progressed) G-CIMP-low. In addition, we identified distinct molecular patterns that distinguish G-CIMP-low recurrent tumor entities from their G-CIMP-high primary counterparts, and defined and validated candidate predictive biomarker signatures that defines risk to G-CIMP-low progression with clinical relevance among primary gliomas harboring IDH-mutant non-codel.

**Table 1.**
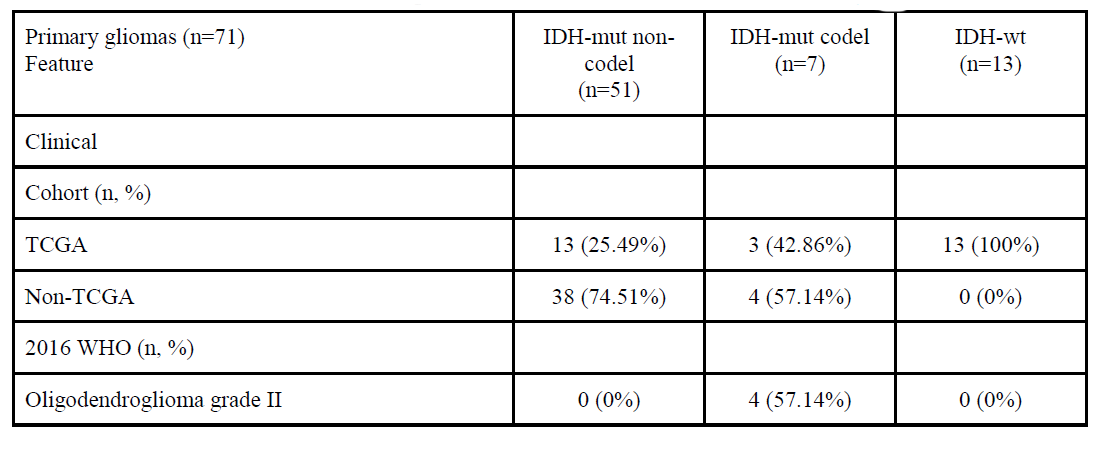
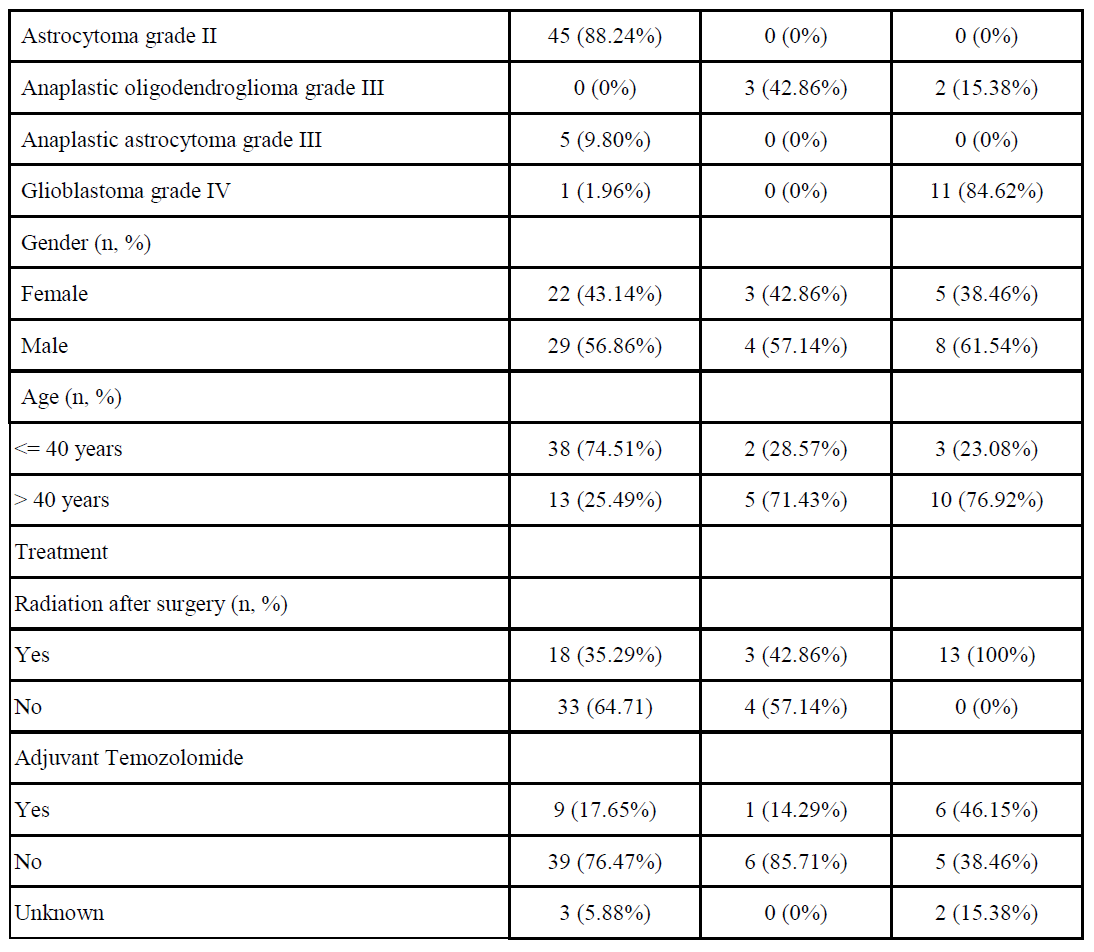
Clinical characteristics of the primary sample cohort arranged by IDH1/1p-19q status Percentages were calculated as a proportion of a total amount of tumor samples in the primary glioma cohort by IDH1/1p-19q status. In cases where more than one tumor fragment per primary surgery were investigated, each case was counted once to avoid overrepresentation of data.

| Primary gliomas (n=71)<br>Feature | IDH-mut non-codel<br>(n=51) | IDH-mut codel<br>(n=7) | IDH-wt<br>(n=13) |
| --- | --- | --- | --- |
| Clinical |  |  |  |
| Cohort (n, %) |  |  |  |
| TCGA | 13 (25.49%) | 3 (42.86%) | 13 (100%) |
| Non-TCGA | 38 (74.51%) | 4 (57.14%) | 0 (0%) |
| 2016 WHO (n, %) |  |  |  |
| Oligodendroglioma grade II | 0 (0%) | 4 (57.14%) | 0 (0%) |
| Astrocytoma grade II | 45 (88.24%) | 0 (0%) | 0 (0%) |
| Anaplastic oligodendroglioma grade III | 0 (0%) | 3 (42.86%) | 2 (15.38%) |
| Anaplastic astrocytoma grade III | 5 (9.80%) | 0 (0%) | 0 (0%) |
| Glioblastoma grade IV | 1 (1.96%) | 0 (0%) | 11 (84.62%) |
| Gender (n, %) |  |  |  |
| Female | 22 (43.14%) | 3 (42.86%) | 5 (38.46%) |
| Male | 29 (56.86%) | 4 (57.14%) | 8 (61.54%) |
| Age (n, %) |  |  |  |
| <= 40 years | 38 (74.51%) | 2 (28.57%) | 3 (23.08%) |
| > 40 years | 13 (25.49%) | 5 (71.43%) | 10 (76.92%) |
| Treatment |  |  |  |
| Radiation after surgery (n, %) |  |  |  |
| Yes | 18 (35.29%) | 3 (42.86%) | 13 (100%) |
| No | 33 (64.71) | 4 (57.14%) | 0 (0%) |
| Adjuvant Temozolomide |  |  |  |
| Yes | 9 (17.65%) | 1 (14.29%) | 6 (46.15%) |
| No | 39 (76.47%) | 6 (85.71%) | 5 (38.46%) |
| Unknown | 3 (5.88%) | 0 (0%) | 2 (15.38%) |

**Table 2.**
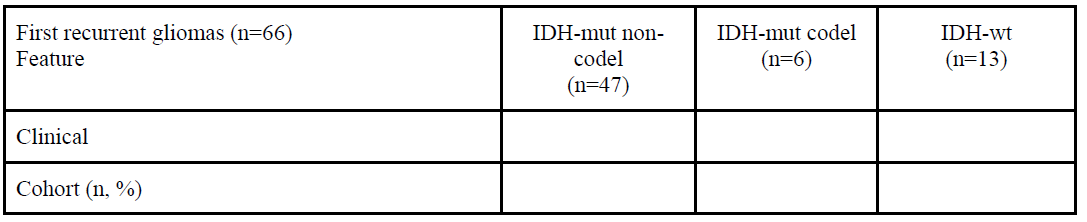
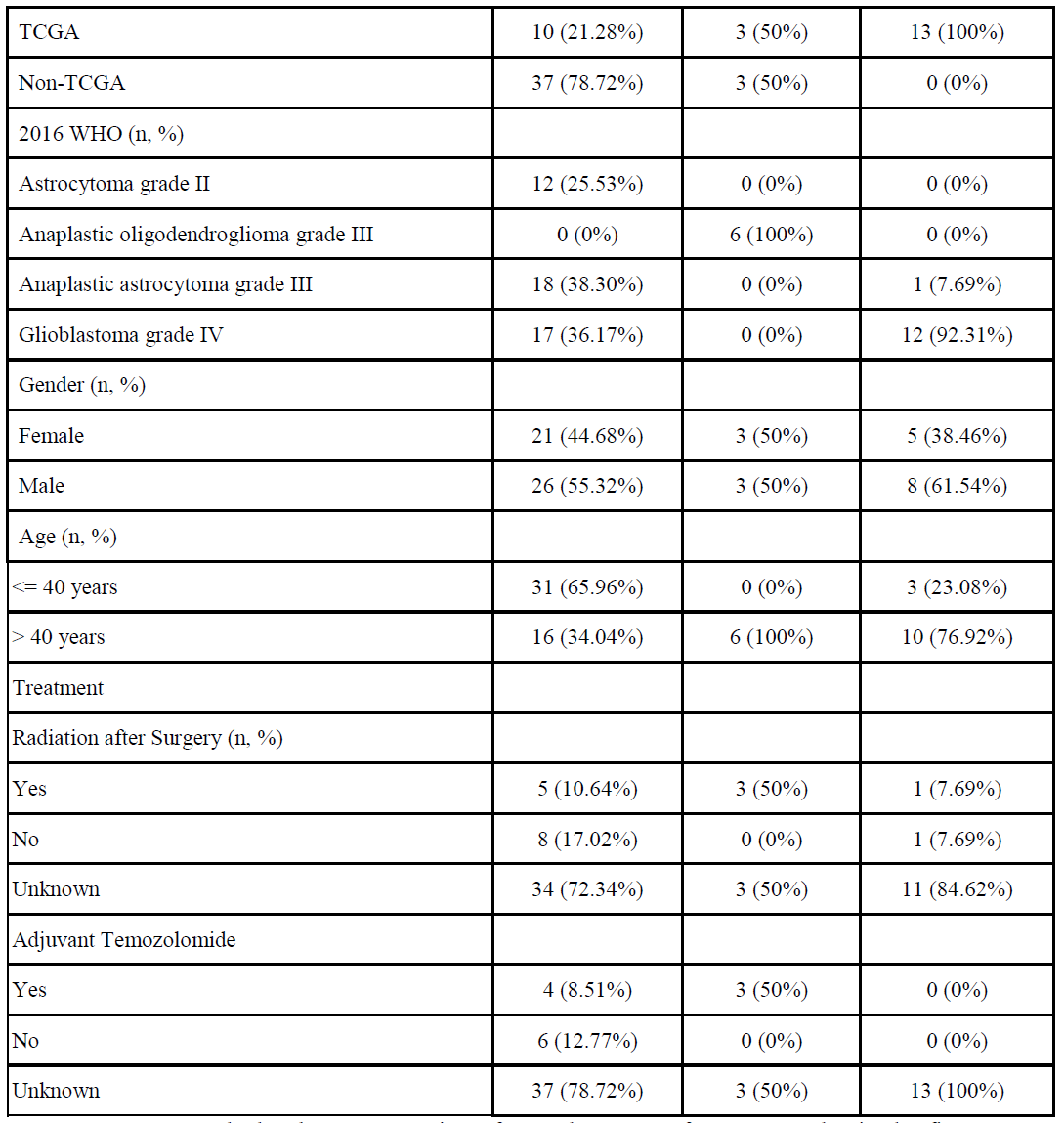
Clinical characteristics of the first recurrent sample cohort arranged by IDH1/1p-19q status Percentages were calculated as a proportion of a total amount of tumor samples in the first recurrent glioma cohort by IDH1/1p-19q status. In cases where more than one tumor fragment per first recurrent surgery were investigated, each case was counted once to avoid overrepresentation of data.

| First recurrent gliomas (n=66)<br>Feature | IDH-mut non-codel<br>(n=47) | IDH-mut codel<br>(n=6) | IDH-wt<br>(n=13) |
| --- | --- | --- | --- |
| Clinical |  |  |  |
| Cohort (n, %) |  |  |  |
| TCGA | 10 (21.28%) | 3 (50%) | 13 (100%) |
| Non-TCGA | 37 (78.72%) | 3 (50%) | 0 (0%) |
| 2016 WHO (n, %) |  |  |  |
| Astrocytoma grade II | 12 (25.53%) | 0 (0%) | 0 (0%) |
| Anaplastic oligodendroglioma grade III | 0 (0%) | 6 (100%) | 0 (0%) |
| Anaplastic astrocytoma grade III | 18 (38.30%) | 0 (0%) | 1 (7.69%) |
| Glioblastoma grade IV | 17 (36.17%) | 0 (0%) | 12 (92.31%) |
| Gender (n, %) |  |  |  |
| Female | 21 (44.68%) | 3 (50%) | 5 (38.46%) |
| Male | 26 (55.32%) | 3 (50%) | 8 (61.54%) |
| Age (n, %) |  |  |  |
| <= 40 years | 31 (65.96%) | 0 (0%) | 3 (23.08%) |
| > 40 years | 16 (34.04%) | 6 (100%) | 10 (76.92%) |
| Treatment |  |  |  |
| Radiation after Surgery (n, %) |  |  |  |
| Yes | 5 (10.64%) | 3 (50%) | 1 (7.69%) |
| No | 8 (17.02%) | 0 (0%) | 1 (7.69%) |
| Unknown | 34 (72.34%) | 3 (50%) | 11 (84.62%) |
| Adjuvant Temozolomide |  |  |  |
| Yes | 4 (8.51%) | 3 (50%) | 0 (0%) |
| No | 6 (12.77%) | 0 (0%) | 0 (0%) |
| Unknown | 37 (78.72%) | 3 (50%) | 13 (100%) |

## RESULTS

### Samples and clinical data

A summary of clinical data features is represented in Tables 1 and 2 and reflects our effort to manually update the available information at the TCGA Biospecimen Core Resource (BCR) combined with previously published datasets (Mazor et al., 2015; Bai et al., 2016) as well as our own cohort with known IDH mutation and 1p-19q codeletion status. The majority of samples were IDH-mutant non-codel at primary (51 out of 71, 71.83%) and first recurrent (47 out of 66, 71.21%) surgery time points. Stratification of histology and grading among the IDH-mutant cases, included astrocytoma grade II as primary (45, 88.24%), anaplastic astrocytoma grade III (18, 38.3%) and glioblastoma grade IV (17, 36.17%) at first recurrence.

### Identification of longitudinal tumors with G-CIMP-high to G-CIMP-low transition

Our group and others have reported on the widespread differences in DNA methylation in adult diffuse primary gliomas (Sturm et al., 2012; Cancer Genome Atlas Research Network et al., 2015; Ceccarelli et al., 2016). We grouped tumors into two IDH-driven macro-clusters eventually leading to the identification of three IDH-mutant specific DNA methylation subtypes (G-CIMP-high, G-CIMP-low, and Codel) and three IDH-wildtype specific DNA methylation subtypes (Classic-like, Mesenchymal-like, and LGm6). Based on the molecular similarity with pilocytic astrocytomas (PA), LGG tumors described as LGm6 pan-glioma methylation subtype were further labeled as PA-like. Additionally, the GBMs falling into this group were best described as LGm6-GBM for their original pan-glioma DNA methylation cluster and tumor grade (Ceccarelli et al., 2016).

TCGA glioma samples not classified in our previously published analysis (n=39; 9 primary and 30 recurrent) in addition to 20 matched primary cases previously included were classified into one of the seven DNA methylation subtypes. To do this we applied a machine learning prediction model (Random Forest [RF]) using our previously defined DNA methylation signatures (Ceccarelli et al., 2016) (IDH-mutant tumor specific (n=1,308), IDH-mutant subtypes (n=163), and 914 IDH-wildtype tumor specific). We extended our analysis by similarly assigning each tumor sample in the non-TCGA longitudinal published cohorts (Mazor et al., 2015) [n=62] and (Bai et al., 2016) [n=48] to one of the DNA methylation subtypes. Additionally, we classified a total of 12 new primary and recurrent glioma samples generated from our own cohort and classified 9 tumor fragments derived from biopsies of 3 different patients (Table S1).

Since IDH mutation status is the dominant molecular feature to classify the cohort into one of the seven DNA methylation subtypes, we integrated an additional set of 1,300 tumor-specific probes that discriminated the pan-glioma cohort into two macro-groups: the LGm1/LGm2/LGm3 DNA methylation macro group harboring *IDH1* or *IDH2* mutations *versus* the LGm4/LGm5/LGm6 DNA methylation macro group comprising glioma samples carrying IDH-wildtype (Ceccarelli et al., 2016). Therefore, the DNA methylomes of 181 longitudinally collected TCGA and non-TCGA gliomas from 74 patients profiled on the Illumina HumanMethylation450 bead arrays (450K) platform were classified according to the CpG probe signatures that define the seven cohesive pan-glioma DNA methylation subtypes described in Ceccarelli et al., 2016. Of the 181 glioma fragments, 122 (67.4%) were classified as G-CIMP-high, 20 (11%) were classified as Codel, 10 (5.5%) were classified as G-CIMP-low, 12 (6.6%) were classified as Mesenchymal-like, 11 (6.1%) were classified as Classic-like, 5 (2.8%) were classified as PA-like, and 1 (0.6%) was classified as LGm6-GBM by supervised RF computational approaches with high specificity and sensitivity (accuracy > 95% on average) (Figures 1A and 1B and Table S1).

**Figure 1.**
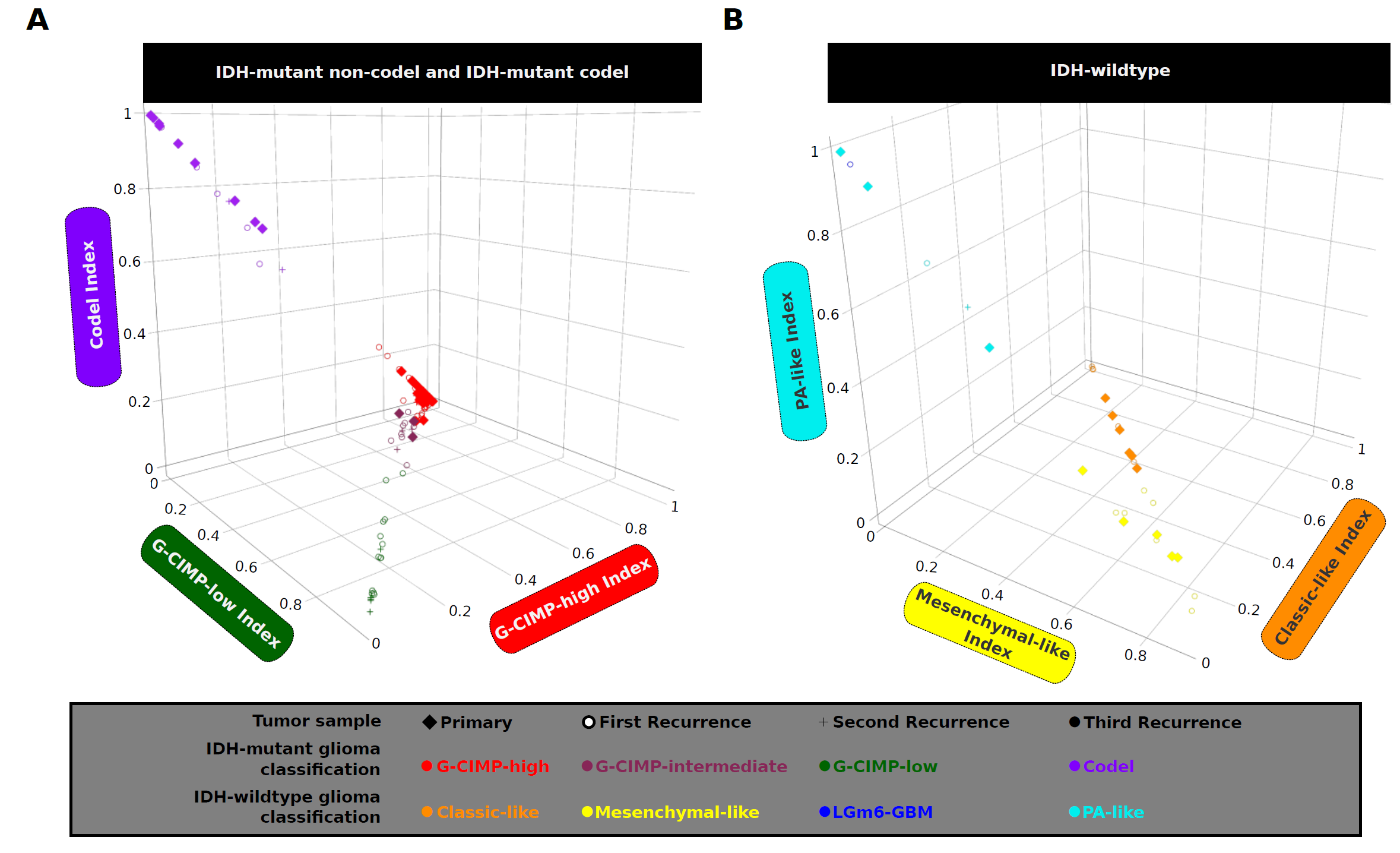
Identification of longitudinal tumors with G-CIMP-high to G-CIMP-low epigenetics shift during recurrence and malignant tumor progression. The methylomes of 181 longitudinally collected TCGA and non-TCGA adult diffuse grade II-III-IV gliomas from 74 patients profiled on the HM450 platform were classified by supervised Random Forest (RF) computational approaches into one of the 7 pan-glioma DNA methylation subtypes using the CpG probe signatures described in Ceccarelli et al., 2016. (A) This three-dimensional (3D) scatter plot using IDH-mutant codel and IDH-mutant non-codel G-CIMP-high and G-CIMP-low indices predicted by RF model shows a distinct set of samples within the IDH-mutant non-codel G-CIMP subgroups exhibiting relatively intermediate DNA methylation profile. These samples have been named G-CIMP-intermediate post-RF assessment. A subset of recurrent glioma projects a G-CIMP-low phenotype while a subset retains their original G-CIMP-high phenotype. (B) (3D) scatter plot using IDH-wildtype PA-like, classic-like, and mesenchymal-like indices predicted by RF shows that IDH-wildtype gliomas did not change significantly in terms of their epigenome profile during disease relapse.

Despite harboring IDH mutation, G-CIMP-low tumors were reported to have lower DNA methylation levels and worse clinical outcome when compared with G-CIMP-high tumors (Ceccarelli et al., 2016)., A three-dimensional (3D) scatter plot using G-CIMP-low and G-CIMP-high indices predicted by RF model (Figure 1A) allowed us to visualize the phenotypic relationships between primary and recurrent G-CIMP-positive tumors, suggesting a distinct set of samples within the IDH-mutant non-codel G-CIMP subgroups that showed relatively intermediate DNA methylation profile at G-CIMP-low index threshold of < 0.5 & ≥ 0.2 and at G-CIMP-high index threshold of ≥ 0.5 & < 0.75. We have named this group of samples as G-CIMP-intermediate post-RF assessment (i.e. n=3 primary, n=7 first recurrent, and n=3 second recurrent tumor fragments derived from 11 distinct patients). G-CIMP-intermediate group can be characterized by modest degree of epigenomic changes trending towards G-CIMP-low (Figures 1A and 2A and Table S1). This may suggest that G-CIMP-intermediate reflects an early stage transition from G-CIMP-high to G-CIMP-low. Notably, we demonstrated a dramatic epigenomic shift towards malignant progression from primary G-CIMP-high to first recurrent G-CIMP-low in seven patients (Figure 2A). Although all G-CIMP-low samples at recurrence are grade IV, not all grade IV tumors transition to G-CIMP-low suggesting that grade may not be the only indicator of G-CIMP-low progression (Figure 2A). We did not observe any significant changes in the IDH-mutant codel and IDH-wildtype glioma subgroups in terms of their epigenome profile towards recurrent disease (Figures S1A and S1B and Table S1). Taken together, epigenomic profiling of longitudinally diffuse gliomas (n=74 patients) confirms and expands our previous hypothesis that G-CIMP-low tumors progress from primary G-CIMP-high tumors (Ceccarelli et al., 2016).

**Figure 2.**
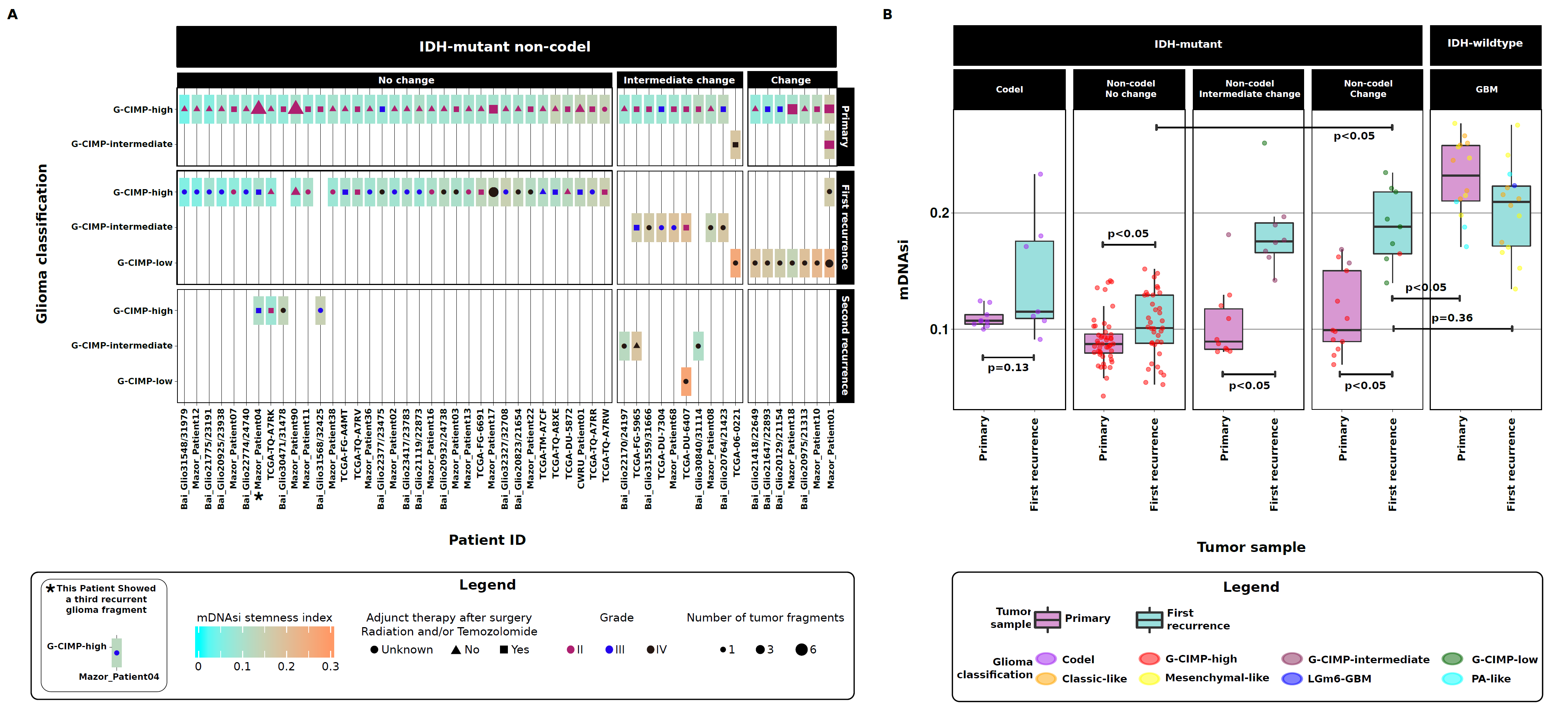
A stem cell-like phenotype defines G-CIMP-low recurrent tumors that resemble IDH-wildtype primary GBMs. An overview of the sample cohort (n=74 patients) across all tissue source sites is shown and highlights the stratification of glioma patients defined by their epigenomic profiles at primary and recurrent surgery time points. (A) A subset within the IDH-mutant non-codel G-CIMP-high subgroup that retains their original epigenomics phenotype at recurrent disease; a subset within the IDH-mutant non-codel macro group manifesting the “G-CIMP-intermediate” DNA methylation profile at primary and/or recurrent stages plus a subset within the IDH-mutant non-codel macro group exhibiting the G-CIMP-low phenotype on second recurrence (these sets of patients are collectively defined as those showing intermediate shifts in their epigenomic profiles); a subset within the IDH-mutant non-codel macro group that significantly shift their DNA methylation patterns from G-CIMP-high at primary diagnosis to G-CIMP-low at first recurrence. Each box represents a patient tumor colored according to its DNA methylation-based stemness index at primary/recurrent diagnosis. A symbol color, size, and shape within each box represents tumor grade, number of tumor fragments, and adjunct therapy (radiation and/or Temozolomide) received after surgery at primary/recurrent diagnosis. (B) G-CIMP-low recurrent tumors possess a high stemness index when compared to their G-CIMP-high primary counterparts and when compared to G-CIMP-high recurrent tumors. On the other hand, the landscape of stemness index in G-CIMP-low recurrent tumors is highly similar to those found in primary IDH-wildtype (GBM) tumors.

### Recurrent G-CIMP-low samples gain stem cell-like phenotype and resemble IDH-wildtype GBMs

G-CIMP-low primary tumors show molecular features associated with a stem cell-like phenotype (SOX binding elements) with worst overall outcome within IDH-mutant non-codels (Ceccarelli et al., 2016). To explore the relationship of stemness, aggressiveness and G-CIMP malignant tumor progression in detail, we performed comprehensive and multi-platform analyses of the stemness signatures across normal cells with distinct degrees of pluripotency and 33 cancer types, totaling more than 10,000 samples. These analyses resulted in the definition of an independent DNA methylation-based stemness index that provide a metric to categorize tumors according to their patterns of similarity in relation to stem cells (manuscript in preparation - Malta et al). The DNA methylation-based stemness index (mDNAsi) is able to recapitulate known features of stemness in distinct cancer types such as the basal breast subtype and the IDH-wildtype mesenchymal-like and classic-like subtypes (data not shown -manuscript in preparation - Malta et al). In this study, when we applied mDNAsi to our entire matched primary and recurrent glioma cohort we were able to define the degree of undifferentiation as a function of glioma progression (Figure 2B). IDH-wildtype primary and first recurrent samples had the highest overall stemness index (medians of 0.23 and 0.2, respectively) compared to the entire IDH-mutant cohort (medians of 0.1 and 0.13, respectively), however the degree of stemness within IDH-wildtype shifted from primary to first recurrence (*p* < 0.05) suggesting IDH-wildtype recurrent samples may be defined by expansion of a resistant clone that is more differentiated yet aggressive in nature, as reported in metastatic melanoma cells (Cheli et al., 2011). Interestingly, G-CIMP-low first recurrent samples showed higher overall stemness index in relation to their primary G-CIMP-high counterparts (*p* < 0.05; medians of 0.19 and 0.1, respectively). When compared to first recurrent G-CIMP-high tumors, the landscape of stemness index in first recurrent G-CIMP-low tumors highly resembles those found in primary IDH-wildtype (GBM) tumors (*p* < 0.05; medians of 0.1 and 0.2, respectively) (Figures 2A and 2B). It is known that IDH-wildtype GBM tumors are more aggressive and can be defined by their de-differentiated state due to a small subpopulation of cancer stem cells (Singh et al., 2004), characterized by self-renewal ability and multi-lineage differentiation potential. In addition, glioma stem cells have been shown to be resistant to adjuvant therapy leading to tumor recurrence (Vescovi et al., 2006; Lathia et al., 2015). Therefore, we defined a subset of tumors at first recurrence that resembles G-CIMP-low primary and they acquire a stem cell-like phenotype upon progression. This suggests that a stem cell-like aggressive tumor behavior exists within a subset of gliomas harboring IDH mutation and intact chromosome 1p and 19q arms, which may contribute to their resistance to adjuvant therapy and relapse as G-CIMP-low phenotype (Figure 2A).

### G-CIMP-high to G-CIMP-low transition is defined by epigenomic changes at known genomic functional elements

We performed a supervised analysis to determine distinct epigenetic changes between the groups defined by a glioma subtype shift (Figures 2A and S1). We did not identify any significant epigenetic difference between 1) primary and recurrent IDH-wildtype gliomas; 2) primary and recurrent Codel gliomas; or 3) primary and recurrent G-CIMP-high tumors. Using a core set of nine cases that significantly shift their DNA methylation patterns from G-CIMP-high at primary diagnosis to G-CIMP-low at first recurrence (Figure 2A and Table S1) we identified 684 differentially hypomethylated CpG probes and 28 differentially hypermethylated probes (FDR < 0.05, difference in mean methylation beta value > 0.5 and < −0.4) associated with G-CIMP-low recurrence (Figure 3A). When we compared these 712 G-CIMP-low signature to non-tumor, normal neuronal and normal glial cells, we observed that the G-CIMP-high (primary and recurrent) tumors are normal-like, contrary to what we found for G-CIMP-low recurrence and grade IV IDH-wildtype (primary and recurrent). Therefore, the 712 G-CIMP-low recurrent CpG signatures were able to stratify IDH-mutant non-codel G-CIMP tumors exhibiting progressed disease and highly aggressive (IDH wildtype-like) phenotypes (Figure 3A). This finding (Figure 3A) combined with our stemness analysis (Figure 2) demonstrate that G-CIMP-low recurrent tumors shared epigenetic characteristics with IDH-wildtype primary GBMs. Although recurrent G-CIMP-low can be classified as grade IV (7/17, 41%), not all grade IV recurrent progressed to a G-CIMP-low phenotype, in fact 35% (6/17) of grade IV at recurrence can be characterized as G-CIMP-high while 24% (4/17) were characterized as G-CIMP-intermediate (Figure 2A). To evaluate if there are differences within grade IV, we performed a supervised DNA methylation analysis between recurrent grade IV G-CIMP-low (n=9) and recurrent grade IV G-CIMP-high (n=6). We observed 350 differentially methylated probes (Wilcoxon rank-sum test, *p* < 0.01 and difference in mean methylation beta value < −0.4 and > 0.5, Figure S2). Collectively, these findings suggest that G-CIMP-high to G-CIMP-low follows an alternative epigenetic roadmap towards disease relapse independent of grade (Figure S2).

**Figure 3.**
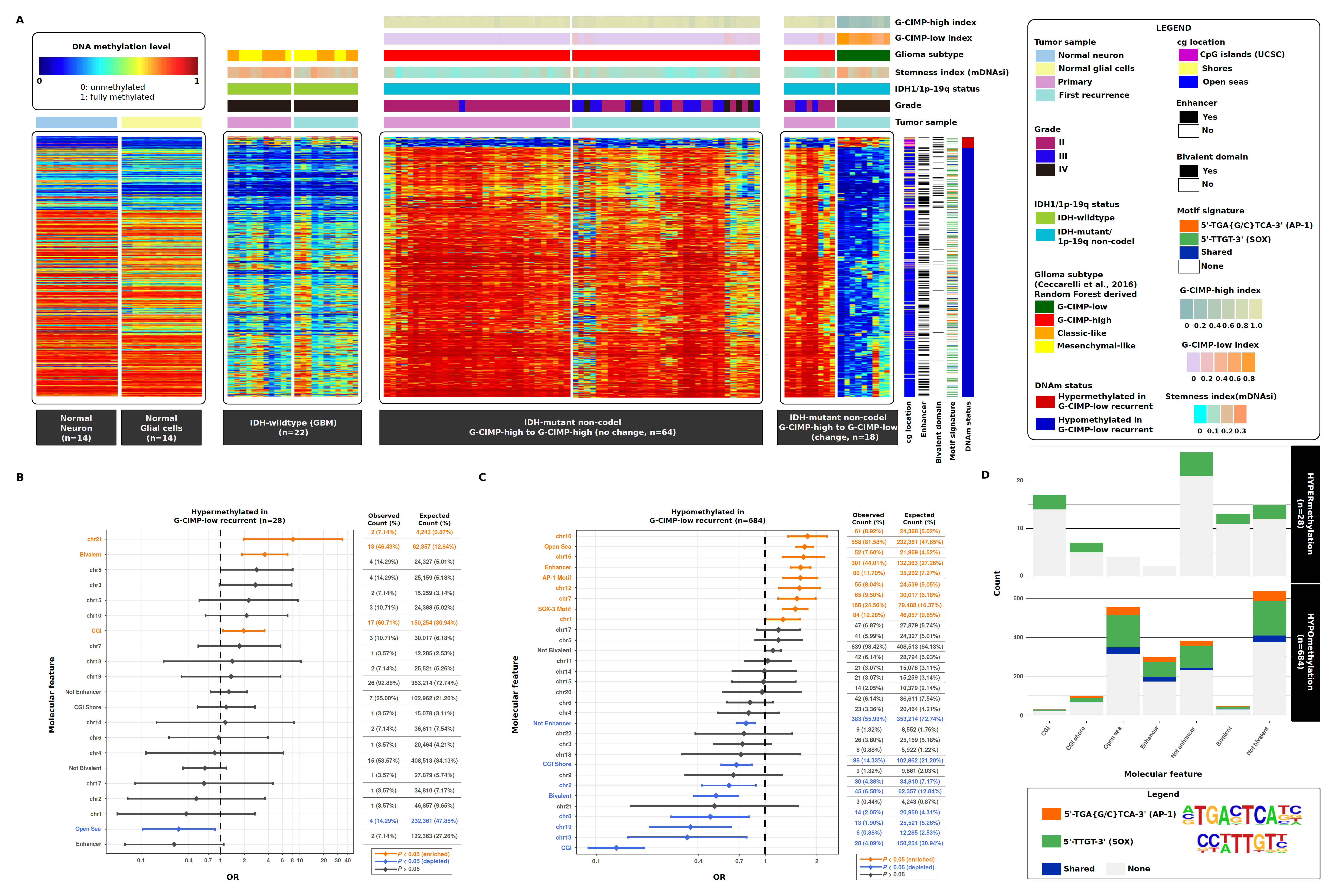
G-CIMP-low recurrent tumors and IDH-wildtype primary GBMs share deregulated DNA methylation events at known genomic functional elements. (A) Heatmap of DNA methylation data. Columns represent non-tumor brain cells (normal neuron cells and normal glial cells, n=28), IDH-wildtype (GBM) (n=22), and IDH-mutant non-codel gliomas (n=64) grouped according to their epigenomic profile at primary and recurrent surgery time points. Rows represent probes identified after supervised analysis between DNA methylation of G-CIMP-high tumors at primary diagnosis and their G-CIMP-low counterparts at first recurrence sorted by hierarchical clustering (n=28 hypermethylated probes and n=684 hypomethylated probes in G-CIMP-low recurrent tumors; FDR < 0.05, difference in mean methylation beta value < −0.4 and > 0.5). Labels on the top and tracks on the right of the heatmap represent clinical and molecular features of interest. The saturation of either color scale reflects the magnitude of the difference on DNA methylation level. (B-C) Odds Ratio (OR) for the frequencies of differentially hypermethylated probes and differentially hypomethylated probes, respectively, that overlap a particular molecular feature relative to the expected genome-wide distribution of 450k probes. (D) *De novo* and known motif scan analyses identified recurring patterns in DNA that are presumed to have sequence binding specific sites for c-JUN/AP1 (5’-TGA{G/C}TCA-3’) and SOX family of transcription factors (5’-TTGT-3’). The molecular features overlapping both motif signatures are shown.

CpG sites exhibiting DNA hypermethylation in G-CIMP-low recurrence were significantly enriched for CpG islands (CGI) (Odds Ratio [OR]=1.96, 95% CI: 1.07-3.58), bivalent chromatin domains (OR=3.61, 95% CI: 1.87-6.98), and chromosome (Chr) 21 (OR=8.17, 95% CI: 1.95-34.32) (Figure 3B, *p* < 0.05 [enriched]). We also observed a depletion of probes positioned within intergenic regions or open seas (OR=0.30, 95% CI: 0.10-0.85) (Figure 3B, *p* < 0.05 [depleted]). Markedly, CpG sites showing DNA hypomethylation in G-CIMP-low recurrence were significantly enriched for open seas (OR=1.70, 95% CI: 1.52-1.91), enhancer elements (OR=1.61, 95% CI: 1.41-1.85), and Chr 1, 7, 10, 12, and 16 (OR>1.0) (Figure 3C, *p* < 0.05 [enriched]). However, we observed a depletion of probes located at CGI (OR=0.13, 95% CI: 0.09-0.19), genomic regions of 2,000 bp upstream and downstream flanking CGI boundaries known as CGI shores (OR=0.67, 95% CI: 0.55-0.83), bivalent chromatin domains (OR=0.51, 95% CI: 0.38-0.69), non-enhancer elements (OR=0.77, 95% CI: 0.68-0.87), and Chr 2, 8, 13, and 19 (OR<0.3) (Figure 3C, *p* < 0.05 [depleted]). Genomic abnormalities pertaining to chromosomes 1, 7, 10, 12, and 19 were documented in gliomas (Brennan et al., 2013; Cancer Genome Atlas Research Network et al., 2015) which may suggest that CpG sites residing on these chromosomes are aberrantly demethylated as a consequence of chromosomal alteration. The majority of CpG sites that underwent a massive DNA demethylation in G-CIMP-low recurring tumors were primarily found within intergenic open sea regions (558 out of 684, 81.58%) (Figures 3A and 3C), a finding consistent with our previous study of primary G-CIMP-high and primary G-CIMP-low tumors (Ceccarelli et al., 2016).

By aggregating chromHMM data from NIH Roadmap Epigenomics Consortium (Roadmap Epigenomics Consortium et al., 2015) with the 712 differentially methylated regions identified in G-CIMP-low progressed tumors, we observe these genomic elements to be functionally relevant in defining differentiated adult tissue phenotype and pluripotency in stem cells (Figure S3). Loss of CpG methylation at these known functional genomic elements associated with normal development and pluripotency defines a possible mechanism of glioma progression which may lead to improved targeted therapy for G-CIMP-low.

### G-CIMP-low recurrent demethylation events are defined by c-JUN/AP-1 and SOX binding elements

DNA methylation signatures of multiple disease-related genes and intergenic regions have recently been related to mortality outcomes (Zhang et al., 2017), providing evidence for the collaborative role of DNA methylation and non-coding functional regions in the modulation of cell phenotypes. For a more functional view of the recurring patterns in hypomethylated DNA that are presumed to have sequence binding specific sites for transcription factors implicated in tumor relapse and progression to G-CIMP-low (n=684 CpG sites), we performed *de novo* and known DNA motif scan analyses. The top ranked *de novo* motif signature, 5’-TGA{G/C}TCA-3’ (geometric test *p*=1e-16, fold enrichment=3.04), corresponded to known motifs associated with the transcription factors JUN-AP1 (geometric test *q*=0, fold enrichment=2.86), FOSL2 (geometric test *q*=0, fold enrichment=2.42), FRA1 (geometric test *q*=0, fold enrichment=2.01), BATF (geometric test *q*=2e-4, fold enrichment=1.72), ATF3 (geometric test *q*=4e-4, fold enrichment=1.67), and AP1 (geometric test *q*=7e-4, fold enrichment=1.60). AP-1 (activating protein-1) is a collective term referring to homodimeric or heterodimeric transcription factors composed of basic region-leucine zipper (bZIP) proteins JUN, FOS or ATF subfamilies. AP-1 is involved in cellular proliferation, transformation, and death (Shaulian and Karin, 2002). We found that AP-1 may significantly bind to probes of demethylated DNA (80 out of 684, 11.70%) in G-CIMP-low progressed cases (OR=1.61, 95% CI: 1.28-2.03) (Figure 3C). From the list of 684 hypomethylated regions, we then extracted those that mapped to the DNA motif signature 5’-TGA{G/C}TCA-3’ (n=87 differentially methylated regions [DMR]). Among them, 87.36% were located in open seas and 59.77% overlapped with enhancers known to define tissue phenotype (76 and 52 DMR, respectively) (Figures 3A and 3D).

Our findings also suggest a motif signature 5’-TTGT-3’, known to be associated with SOX transcription factor family members, as significantly enriched: SOX3 (geometric test *q*=0, fold enrichment=1.49), SOX6 (geometric test *q*=0, fold enrichment=1.47), SOX2 (geometric test *q*=1e-4, fold enrichment=1.67), SOX10 (geometric test *q*=7e-4, fold enrichment=1.39), SOX4 (geometric test *q*=2.2e-3, fold enrichment=1.55), and SOX15 (geometric test *q*=2.2e-3, fold enrichment=1.47). We observed that SOX transcription factors collectively may bind 226 differentially hypomethylated regions, most of them located in intergenic open sea regions (200 out of 226, 88.50%) (Figures 3A and 3D). The enrichment of SOX-related motifs in recurrent/progressed G-CIMP-low cases is in line with our published findings in a cohort of primary G-CIMP-high and primary G-CIMP-low samples (Ceccarelli et al., 2016) and with our observation that these sites are enriched for functional genomic elements. Additionally, a set of 5 differentially hypermethylated regions, mostly located in CGI, showed DNA binding sites for the SOX-related motif signature (Figures 3A and 3D). Thirty-five out of 684 hypomethylated regions (5.12%) shared both 5’-TGA{G/C}TCA-3’ and 5’-TTGT-3’ motif signatures (Figure 3D). Therefore, the above results suggest that DNA methylation changes on the 712 CpGs deregulated in G-CIMP-low recurrence can alter functional DNA binding sites recognized by c-JUN/AP-1 and SOX elements contributing to G-CIMP-low malignant tumor progression (compilation of results shown in Figure 3).

### RNA fusions and genomic rearrangements abnormalities associated with G-CIMP progression

We investigated RNA fusions and genomic rearrangements associated with G-CIMP progression. Since we did not have RNA fusion and genomic data for all cases in this study, in particular within G-CIMP-low, we focused our attention on the G-CIMP-intermediate group. We observed G-CIMP-intermediate to be likely an early stage transition from G-CIMP-high to G-CIMP-low as observed by the overall similarity between stemness index and DNA demethylation state at recurrence (Figures 1 and 2). Interestingly, G-CIMP-intermediate at recurrence reflects major genomic rearrangements and loss of cell cycle related genes (*CDKN2A*, *CDKN1B*) compared to G-CIMP-high at primary and recurrence (Figure S4). Given the limited number of samples, specific RNA fusions driving G-CIMP-intermediate was not observed. However, we did observe G-CIMP-high tumors which retained their epigenomic signature (e.g. G-CIMP-high at primary and recurrence) tend to recur with similar rearrangement events while tumors with an epigenomic shift (G-CIMP-high to intermediate) had an increase in number of differing rearrangement events. This suggests a subclonal evolution of tumor cells emerging from primary to recurrence for G-CIMP-intermediate and likely G-CIMP-low (Figure S4) that may be resistant to therapy (Figure 2A).

### Predictive biomarker signatures can predict risk to G-CIMP-low progression at primary diagnosis

Our study highlights that G-CIMP-low recurrent tumor entities resemble tumors which belonged to the IDH-wildtype GBM group known to exhibit an aggressive phenotype (Figures 2 and 3). Currently, glioma relapse and malignant tumor progression are unpredictable by histomorphology and grading scheme (Sanai et al., 2011; Louis et al., 2016). To test whether G-CIMP-high to G-CIMP-low progression can be predicted from primary G-CIMP-high gliomas, we performed supervised analysis between DNA methylation of primary G-CIMP-high tumors progressing to the G-CIMP-low phenotype and primary G-CIMP-high tumors that retain the G-CIMP-high epigenetic profiling through glioma recurrence (Wilcoxon rank-sum test, *p* < 0.05, absolute difference in mean methylation beta value > 0.2). We defined a set of candidate biomarker signatures composed of seven hypomethylated CpG sites in primary G-CIMP-high tumors that shift their epigenomic profile upon disease relapse (Figure 4A).

**Figure 4.**
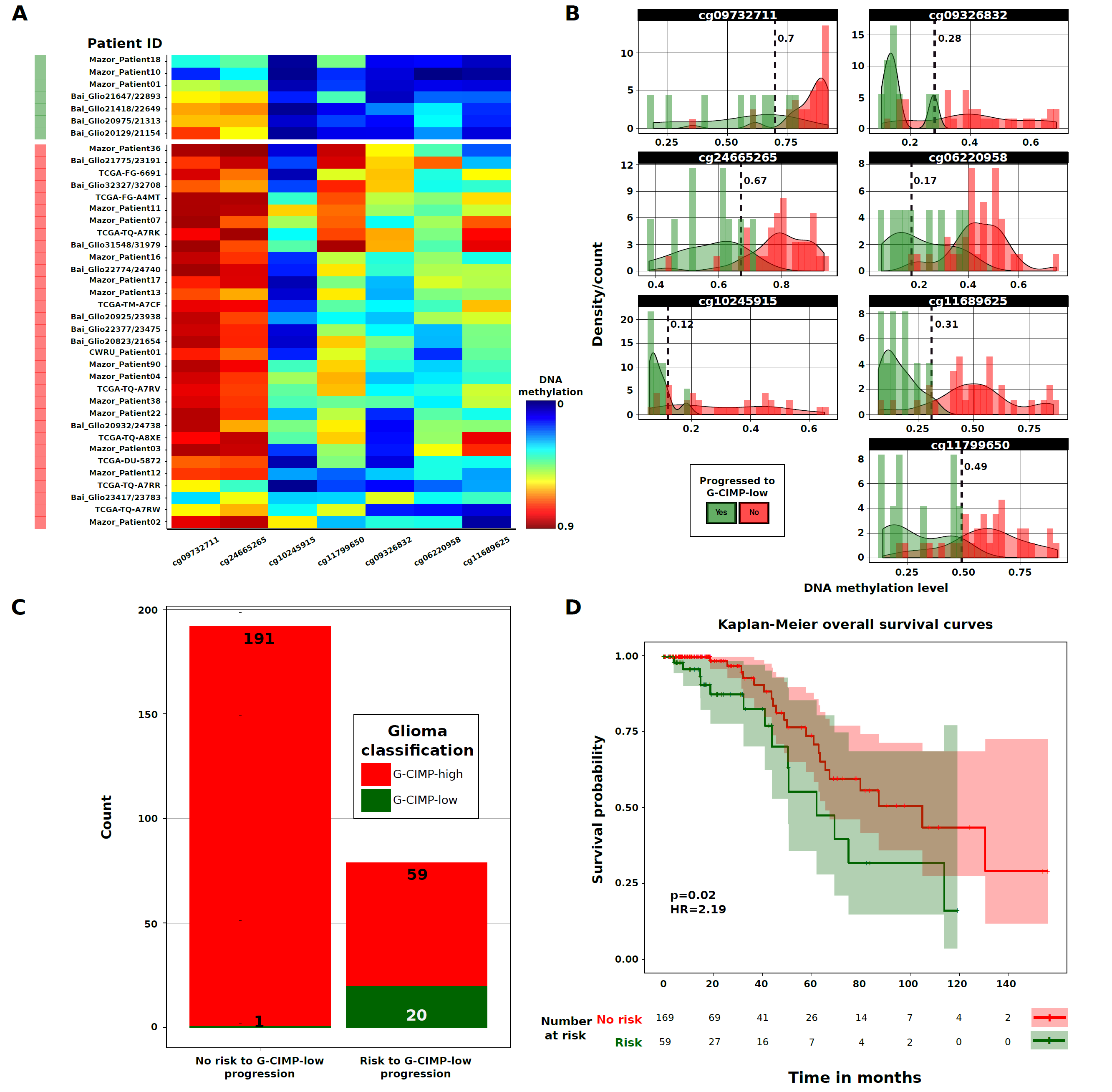
Clinical application of malignant progression to G-CIMP-low. (A) Heatmap of DNA methylation data. Rows represent G-CIMP-high primary cases that progress to G-CIMP-low at recurrent disease (labeled in green) and G-CIMP-high primary cases that retain their original G-CIMP-high phenotype at recurrence (labeled in red). Columns represent the candidate predictive biomarkers identified after supervised analysis of DNA methylation between the two groups mentioned above and are sorted by hierarchical clustering (n=7; unadjusted *p* < 0.05, absolute difference in mean methylation beta value > 0.2). The saturation of either color (scale from blue to red) reflects the magnitude of the difference on DNA methylation level. (B) Beta value thresholds that more specifically distinguish the primary glioma cases that will progress to G-CIMP-low phenotype from those primary glioma cases that will relapse without progression to G-CIMP-low are represented and were used to dichotomize the DNA methylation data in an independent validation cohort (n=271). (C-D) Predictive biomarkers of G-CIMP-low progression correlate with patient outcome.

Next, we sought to determine the usefulness of our biomarkers to predict samples at time of initial surgical diagnosis will progress to the G-CIMP-low phenotype. Toward this aim, we dichotomized the data using beta value thresholds which more specifically distinguish the primary glioma cases that will relapse as a progressed G-CIMP-low disease from those primary glioma cases that will relapse without progression to G-CIMP-low. The beta value cutoff for each CpG probe was as follows: cg09732711 (0.7), cg09326832 (0.28), cg24665265 (0.67), cg06220958 (0.17),cg10245915 (0.12), cg11689625 (0.31), and cg11799650 (0.49) (Fisher’s exact test, FDR=0.03, prognostication value and FDR were assigned at n ≥ 5 probes) (Figure 4B). We then investigated and validated the predictive value of these epigenetic biomarker signatures in an independent cohort of 271 TCGA and non-TCGA primary gliomas previously classified in Ceccarelli et al., 2016 as IDH-mutant non-codel G-CIMP-high (n=250) and IDH-mutant non-codel G-CIMP-low (n=21). These 271 primary glioma samples were obtained from published datasets (Sturm et al., 2012; Turcan et al., 2012; Mur et al., 2013; Ceccarelli et al., 2016). We found that the candidate biomarker signatures identified here successfully predicted 29% of tumors (79 out of 271) belonging to the “risk group,” including 95% (20 out of 21) previously identified G-CIMP-low primary tumors (Figure 4C), with clinical relevance on patient overall survival (log-rank *p* value =0.02, hazard ratio [HR] = 2.19) (Figure 4D). These results provide insights into the tumorigenic events that drive G-CIMP progression, with opportunity for further targeted therapy exploitation, as well as a clinical inclusion into clinical trials design to impede or prevent tumor progression to G-CIMP-low, a known aggressive tumor phenotype associated with IDH-mutant non-codel gliomas.

## DISCUSSION

We describe the landscape of epigenomic alterations in adult diffuse G-CIMP tumors through analysis of 181 paired primary and recurrent gliomas from 74 patients. Although recent reports have highlighted pronounced epigenetic differences between primary low-grade gliomas and recurrent high-grade gliomas, these studies have grouped tumors by either grade or genomic alterations. In our study, we took a more holistic approach guided by our recent findings that the current WHO 2016 classification can be further divided by epigenomic subtypes that are prognostically advantageous over both IDH mutation status and grade.

By applying the same machine learning algorithm, which we applied to validate our epigenomic subtype (Ceccarelli et al., 2016), we observed that all gliomas retain their genomic subtype (IDH status). Although a majority of the glioma primary and recurrent matched samples retained their epigenomic subtype, a subset of patients’ tumors showed distinct widespread DNA methylation signatures, especially at intergenic regions, that were strongly reflective of (epi)genomic and stemness signatures of IDH-wildtype primary GBMs. Interestingly, these epigenetically distinct tumors at recurrence were classified as G-CIMP-low, a subtype of G-CIMP IDH-mutant non-codels that was shown to be more aggressive than their counterparts (G-CIMP-high IDH-mutant non-codels). In our study we observed 100% of G-CIMP-low at recurrence as grade IV tumors, however, not all grade IV gliomas resemble G-CIMP-low, suggesting that grade may not be the only determinant of G-CIMP-low identification and evolution in this subset of IDH-mutant 1p19q non-codels.

Primary G-CIMP-low tumors are more demethylated at genomic sites associated with intergenic elements defined as enhancers and generally harbor distinct DNA signature or motifs commonly associated with SOX transcription factor binding (Ceccarelli et al., 2016). Similarly, G-CIMP-low recurrent tumors display very similar epigenomic alterations to their primary counterparts (Figure S5) with the addition of c-JUN/AP-1 transcription factor binding motif. Interestingly, a recent study has shown that c-JUN N-terminal phosphorylation regulates the *DNMT1* promoter leading to DNA hypermethylation that is similar to the G-CIMP phenotype in LGGs and proneural GBMs, and correlates with downregulation of mesenchymal-related genes, reduced cell migration and invasiveness (Heiland et al., 2017). This may imply that DNA methylation loss associated with G-CIMP-low recurrence reflects chromatin remodeling events initiated by the crosstalk between c-JUN/AP1 and DNMT1. DNA methylation plasticity captured at least two additional molecular mechanisms of oncogenesis for G-CIMP tumors than either histologic grading and the recently described IDH mutation-driven G-CIMP pattern.

Despite limited genomic and transcriptomic data for this study, we did observe striking differences in gains of distinct genomic alterations and fusions associated with a subset of progressively altered G-CIMP-intermediate tumors compared to G-CIMP-high. Cell cycle related genes were also observed to be deleted or altered among the primary tumors that develop G-CIMP-intermediate phenotype. These included genetic alterations in the *CDKN2A* and *CDKN2B* cell cycle genes, members of the RB (retinoblastoma) and AKT-mTOR (mammalian target of rapamycin) pathways, which were reported previously in recurrent gliomas (Mazor et al., 2015; Bai et al., 2016). Despite the fact that driver mutations in these pathways and the hypermutator phenotype can emerge at disease relapse after adjunct chemotherapy of primary tumors with TMZ (Johnson et al., 2014), we identified convergent genetic alterations in primary G-CIMP-low tumors (Ceccarelli et al., 2016) (Figure S6). This suggests that the aforementioned molecular signatures reflect genomic abnormalities that underlie glioma development and progression in a subset of patients harboring the G-CIMP-low phenotype.

The stratification of G-CIMP-high patients into two macro groups based on molecular changes acquired at first recurrence allowed us to define potential predictive biomarkers with a significant patient survival correlation. Refined prediction of patients harboring primary disease anticipated to have a more aggressive course despite IDH mutation and progression to G-CIMP-low phenotype, as illustrated by this finding, has clinical implications. Such markers that define patient progression at primary diagnosis, could potentially allow one to design *in vitro* and patient derived xenograph models from these fresh tissues to study and evaluate the functional characterization and mechanisms by which G-CIMP-low evolves from G-CIMP-high. Longitudinal monitoring of G-CIMP phenotypes provides insight into the roadmap of glioma progression independent of grade or IDH status, which may allow for exploration of therapeutic targets to disrupt progression to G-CIMP-low and refined clinical trial designs to determine optimal management of patients at risk for progression to G-CIMP-low.

## AUTHOR CONTRIBUTIONS

Conceptualization, H.N. and J.B-S.; Methodology, C.F.S., H.N., T.S.S., T.M.M., A.S., O.M., and S.S.;Validation, C.F.S.; Formal analysis, C.F.S., O.M., T.M.M., and A.S.; Investigation, C.F.S. and H.N.; Resources: C.F.S., H.N., J.B-S., L.S., D.T., C.C., T.M.M., A.S., P.W.L., M.W., A.I., L.P., J.Z., J.S., T.S.S., T.M., W.A.F., K.M., A.C., Z.S., and S.K.; Data curation, C.F.S., H.N., L.S., and J.B-S.; Predictive Biomarkers: C.F.S. and H.N.; Writing – original draft: C.F.S. and H.N.; Visualization: C.F.S. and H.N.; Project administration: H.N.

## ACKNOWLEDGMENTS

The authors are grateful to patients who contributed to this study. H.N., C.F.S., T.S.S., and T.M.M., were supported in part by São Paulo Research Foundation Grants (FAPESP) 2016/15485-8, 2014/08321-3, 2015/07925-5, 2016/01975-3, 2014/02245-3, 2016/12329-5, 2016/06488-3, and institutional grants from Department of Neurosurgery at Henry Ford Hospital. JB-S and LS were supported in part by the Case Comprehensive Cancer Center Grant (NIH/NCI 5P30CA043703 and HHSN261201000057C). MW was supported by Foundation for Polish Science (Welcome grant 2010-3/3). We would like to thank Susan MacPhee for reviewing the draft manuscript. We also like to thank Drs Joseph Costello, Tali Mazor, Murat Gunel and Hanwen Bai for assisting with updates to the clinical and molecular data from their cohorts. The authors report no conflict of interest exists.

## EXPERIMENTAL PROCEDURES

### 1. Biospecimens

Specimens were obtained from patients with appropriate consent from institutional tissue source sites review boards. Molecular data from longitudinally gliomas were generated from our own cohort and obtained from The Cancer Genome Atlas (TCGA) collections and from previously published datasets (Mazor et al., 2015; Bai et al., 2016). Molecular data from normal brain tissue and from primary gliomas were obtained from TCGA and previously published datasets (Sturm et al., 2012; Turcan et al., 2012; Guintivano et al., 2013; Mur et al., 2013; Ceccarelli et al., 2016). Sample ID from our own cohort and published cohorts as well as tissue source sites from TCGA and published data are listed in Table S1. As previously reported (Cancer Genome Atlas Research Network, 2008; Mazor et al., 2015; Bai et al., 2016), biospecimens were collected from patients diagnosed with primary and recurrent LGG and GBM undergoing surgical resection and were either snap frozen in liquid nitrogen or formalin-fixed and paraffin-embedded (FFPE). Biomolecule analyte extraction was conducted in samples after pathology quality control metrics to classify and grade and to estimate the percentage of neoplastic cells in the tumor tissues. In cases where more than one tumor fragment were investigated, each sample was an independent, geographically distinct piece derived from different stages of the same surgery (Cancer Genome Atlas Research Network, 2008; Mazor et al., 2015; Bai et al., 2016).

### 2. Molecular and clinical covariates

Molecular data elements comprise *IDH1* and 1p-19q status. All TCGA cases were manually followed up by the individual tissue source sites and represents a complete update of new cases as well as cases from the previous publication (Brennan et al., 2013; Ceccarelli et al., 2016). Clinical covariates comprise grade, histology, gender, age at primary and recurrent tumor diagnoses, radiation therapy, chemotherapy based on Temozolomide (TMZ), overall survival, and survival since first recurrence. Survival since first recurrence was defined as follows: (patient overall survival) -(time elapsed between primary and first recurrent tumor surgeries) (Table S1). Clinical data were updated as of September 2015 by contacting the individual tissue source sites, in case of TCGA, and as of January 2016 and April 2016 by previously published reports in case of Mazor et al. 2015 and Bai et al. 2016. Survival curves were estimated using the Kaplan-Meier method and Cox proportional hazards analysis.

### 3. DNA isolation and Illumina human methylation 450K (HM450) array

Primary and recurrent samples from CWRU_Patient01, CWRU_Patient02, and CWRU_Patient03 were obtained from Case Western Reserve University. Recurrent samples from previously deposited primary matched TCGA samples (i.e. TCGA-TQ-A7RR, TCGA-TQ-A7RW and TCGA-FG-6691) were also included in this study to expand the total number of available glioma primary and recurrent pairs. In the case of the TCGA IDH-wildtype GBM recurrent cohort (n=12), the initial primary DNA was profiled using the GoldenGate I and II platforms (∼6,000 CpGs) (Noushmehr et al., 2010; Brennan et al., 2013) while the matched recurrent DNA was profiled using the Infinium HumanMethylation450 bead arrays platform (also commonly referred to as 450K). In order to effectively analyze the epigenomics across all available glioma recurrent samples for this study, we generated 450K Illumina DNA methylation for these primary TCGA IDH-wildtype GBMs (n=12), approved by the TCGA director, Dr. Jean Zenklusen (Table S2). Genomic DNA from the gliomas (n=24 samples) was extracted and purified using AllPrep(r) DNA/RNA/miRNA Universal kit (QIAGEN), following the manufacturer’s instructions. Only DNA samples with an absorbance A260/A280 nm ratio ≥1.8 were used for microarray hybridization. Sodium bisulfite conversion of the genomic DNA was done with EZ DNA methylation Kit (ZymoResearch, USA) following the manufacturer’s guidelines. Briefly, purified genomic DNA (1µg) was bisulfite converted and processed on Infinium HumanMethylation450 bead arrays (Illumina Inc.), as described previously (Ceccarelli et al., 2016). Raw 450K DNA methylation data (samples generated for the purpose of this study) is available through Gene Expression Omnibus (GEO accession number xxxx). All other raw data are available through Genomics Data Commons (in the case of TCGA and the data is accessible via TCGAbiolinks (Colaprico et al., 2016) or described in previous studies (Mazor et al., 2015; Bai et al., 2016).

### 4. Illumina 450K array preprocessing

For level 1 TCGA “Illumina HumanMethylation450” data acquisition (version 12 for LGG and version 6 for GBM) we used the Bioconductor package TCGAbiolinks version 1.1.12 (Colaprico et al., 2016). A total of 59 matched primary and recurrent gliomas, including 35 LGG and 24 GBM samples were profiled through a 450K array platform which interrogates DNA methylation levels of 485,577 human CpG sites. In addition to the TCGA data, we obtained a published dataset of 62 (Mazor et al., 2015) and a dataset of 48 (Bai et al., 2016) longitudinally collected gliomas (complete list of samples and their respective ID is available in Table S1). Probe-level signals for individual CpG sites (raw IDAT files) were subjected to background correction, global dye-bias normalization, calculation of DNA methylation level, and detection p-values (Triche et al., 2013) using the Bioconductor package methylumi version 2.16.0. The DNA methylation level for each locus is measured as a beta-value score (б = (M/(M+U)) in which M and U represent the mean methylated and the mean unmethylated signal intensities for each locus, respectively. Beta-values score range from zero to one with scores of zero indicating complete unmethylated DNA and scores of one indicating complete methylated DNA. A p-value detection of each data point is used to compare the signal intensity difference between the analytical CpG sites and a set of negative control CpG sites represented in the array. A p-value of greater than 0.05 is statistically insignificantly different from the background and is masked as “NA”. Probes that are designed for sequences with known germline polymorphisms (Illumina supplementary SNP list version 2, downloaded 12 December, 2016) and the X and Y chromosomes were filtered out before supervised analysis (466,835 probes remained following filtering).

### 5. Annotation of regulatory genomic elements interrogated by 450K array

The genomic location of 450K probes was divided into those within CpG islands (CGI), in CGI shores, or in open seas (with and without overlapping gene bodies) using UCSC genome table browser (hg19) CpG islands genomic annotation. UCSC adopts the following criteria to define CGI: GC content of at least 50%, length greater than 200 bp, and ratio of observed number of CG dinucleotides to the expected number of G and C in the DNA segment greater than 0.6 (genome.ucsc.edu). We annotated CGI shores as genomic regions of 2,000 bp upstream and downstream flanking CGI boundaries, and open seas as genomic regions located neither in CGI nor in CGI shores. Accordingly, 31% of 450K probes are located within CGI, 21% in CGI shores, and 48% in open seas. 132,363 450K probes show overlapping with candidate enhancers that were previously described by Yao et al., 2015. Briefly, these functional genomic elements are distal 2,000 bp away from transcription start sites (TSS), DNA hypomethylated in tumors when compared with normal controls, correlated to cancer-specific transcription factors, and overlap three enhancer databases: 1) REMC (Roadmap Epigenomics Mapping Consortium), 2) ENCODE (chromHMM for 98 tissues or cell lines), and 3) FANTOM5 (enhancers defined by eRNAs for 400 distinct cell types (Yao et al., 2015). An annotation of 450K bivalent chromatin domains in human embryonic stem cells (hESCs) was obtained from a published dataset. Accordingly, bivalent probes are mostly located in CGI-rich promoters and possess molecular features associated with depletion of retroelements (LINE, SINE, and LTR) in the genomic region directly bordering the TSS, enrichment of H3K4me3 and H3K27me3 histone modifications (minimum overlapping size of 1 Kb), and occupancy by PRC2 and PolII complexes (Court and Arnaud, 2017).

### 6. Classification of longitudinal IDH-wildtype and IDH-mutant glioma samples based on DNA methylation subtypes

Longitudinal glioma samples were classified as either IDH-wildtype (Classic-like, Mesenchymal-like, LGm6-GBM, and PA-like) or IDH-mutant (Codel, G-CIMP-high, and G-CIMP-low) DNA methylation subtypes using the CpG methylation signatures previously defined by our group (Ceccarelli et al.,2016) (tcga-data.nci.nih.gov/docs/publications/lgggbm_2016/PanGlioma_MethylationSignatures.xlsx). We evaluated the performance of the Random Forest (RF) machine learning prediction model by training on a random set of either 80% of 430 IDH-wildtype samples or 80% of 448 IDH-mutant adult diffuse primary LGG-GBM TCGA samples (tcga-data.nci.nih.gov/docs/publications/lgggbm_2016/LGG.GBM.meth.txt), and then evaluated the performance on the remaining 20% of glioma samples. Given the high specificity and sensitivity of our model (accuracy > 95% on average), we tested the prediction model on the primary and recurrent gliomas (n=181 tumor fragments) to classify them according to the seven DNA methylation glioma subtypes using the RF approach and the R packages caret and randomForest. RF probability indices are provided in Table S1.

### 7. Supervised analysis of DNA methylation

We used the Wilcoxon rank-sum test followed by multiple testing using the Benjamini & Hochberg (BH) method for false discovery rate (FDR) estimation (Benjamini and Hochberg, 1995) to identify differentially methylated sites between two groups of study. The 712 probes (Table S3) were defined by comparing a core set of nine cases that significantly shift their DNA methylation patterns from G-CIMP-high at primary diagnosis to G-CIMP-low at first recurrence, using the following criteria: FDR < 0.05 (adjusted *p*) and difference in mean methylation beta value < −0.4 and > 0.5. The 7 probes that define prediction were defined by comparing a core set of 9 primary G-CIMP-high tumors progressing to the G-CIMP-low phenotype and 32 primary G-CIMP-high tumors that retained the G-CIMP-high epigenetic profiling through glioma recurrence. We used the following criteria: *p* < 0.05 (unadjusted) and absolute difference in mean methylation beta value > 0.2.

### 8. Motif discovery

*De novo* and known motif discovery analyses were conducted using Hypergeometric Optimization of Motif EnRichment (HOMER) version 4.9 (Heinz et al.,2010; homer.salk.edu). The perl script findMotifGenome.pl was used to search for conserved DNA binding sequences (8-20 bp motifs) associated with genomic differentially methylated regions (DMR) (adjusted *p* < 0.05), using the following criteria: hg19 genome assembly, 200 bp upstream and downstream flanking each target CpG site, and expected genome-wide distribution of 450K probes as background. Raw outputs from HOMER can be found on the publication portal accompanying this publication.

### 9. Chromatin state (chromHMM) analysis

ChromHMM data, which define 18 distinct chromatin states by combining 6 histone markers (i.e. H3K27ac, H3K4me1, H3K4me3, H3K36me3, H3K27me3, and H3K9me3) across 98 reference human epigenomes were downloaded from the NIH Roadmap Epigenomics Consortium (Roadmap Epigenomics Consortium et al., 2015). Data shown in Figure S3 reflect the chromatin states in somatic cells and stem cells for the genomic regions that map to the 28 probes defined as differentially hypermethylated and 684 probes defined as differentially hypomethylated in G-CIMP-low recurrent cases in relation to their G-CIMP-high primary counterparts.

### 10. Fusion transcript detection and DNA rearrangement analysis

We investigated fusion transcript events and DNA rearrangements in a total of 30 matched primary and recurrent TCGA LGG samples derived from 13 patients with available RNA sequencing and whole-genome sequencing data for the primary tumor and at least one recurrent sample. RNA sequencing reads were analyzed using deFuse package version 0.6.0 (McPherson et al., 2011). Candidate fusions were filtered based on the following deFuse parameters:

- `Splitr_count ≥ 5 (5 or more split reads supporting the fusion)`
- `Span_count ≥ 10 (10 or more spanning reads supporting the fusion)`
- Read_through ∼ “N” (fusion is not a readthrough)
- Adjacent ∼ “N” (fusion does not involve adjacent genes)
- Altsplice ∼ “N” (fusion can not be explained by alternative splicing)
- Min_map_count=1 (at least one spanning read supporting the fusion is uniquely mapped)
- ORF ∼ “Y” (fusion preserves the open reading frame)

The deFuse fusion predictions were manually reviewed using blat analysis (Kent, 2002) of the breakpoint sequence in the UCSC Genome Browser (Kent et al., 2002). Whole-genome DNA rearrangements were identified using the BamBam tool (Sanborn et al., 2013), following the standard recommendation posted on the Five3 Genomics website (dna.five3genomics.com). Briefly, variants with at least 6 reads with an average mapping quality > 30 were ranked as follows: +2 if read support > 15; +4 if split reads are found; +5 if ORF is preserved in a fusion gene; +4 if “Near fusion” (correct orientation but improper phase); +2 if deletion could cause a loss of any part of a gene; +1 if variant interrupts a gene. deFuse and BamBam predictions were combined to define the genomic events identified by both methods. Genomic rearrangements for each of the 13 patients with primary and recurrent tumor samples is plotted as circos plots (Figure S4). A summary of manually curated DNA and RNA rearrangements for each patient is available on the publication portal accompanying this publication.

### 11 Statistics

Data visualization and statistical analysis were performed using R software packages (http://www.r-project.org) and Bioconductor (Gentleman et al., 2004). All R codes used for this study are available as an Rmarkdown document: github.com/BioinformaticsFMRP/SouzaGCIMPRecurrent.

## SUPPLEMENTARY FIGURES

**Supplementary Figure 1.** Schematic representation of longitudinal tumors carrying (A) IDH-mutant 1p-19q codel and (B) IDH-wildtype genotypes. Each box represents a patient tumor colored according to its DNA methylation-based stemness index at primary/recurrent diagnosis. A symbol color, size, and shape within each box represents tumor grade, number of tumor fragments, and adjunct therapy (radiation and/or Temozolomide) received after surgery at primary/recurrent diagnosis.

**Supplementary Figure 2.** Differentially methylated CpG probes between grade IV first recurrent G-CIMP-low tumors and grade IV first recurrent G-CIMP-high tumors supports the evidence that G-CIMP-low tumor progression occurs independent of grade (n=26 hypermethylated probes and n=324 hypomethylated probes in grade IV G-CIMP-low recurrent tumors; FDR < 0.01, difference in mean methylation beta value < −0.4 and > 0.5).

**Supplementary Figure 3.** Integrative analysis of 98 human epigenomes using Roadmap chromHMM datasets. Chromatin states defined by dynamics of epigenomics marks, profiled by the NIH Roadmap Epigenomics Consortium, were integrated to the 712 differentially methylated regions that discriminate G-CIMP-low recurrent from G-CIMP-high primary with the aim of inferring potential genomic regulatory elements overlapping our set of (A-C) hypermethylated probes (n=28) and (D-F) hypomethylated probes (n=684) across 98 reference epigenomes including normal brain and stem cells.

**Supplementary Figure 4.** Chromosomal rearrangements across (A) IDH-mutant non-codel that did not change significantly in terms of their epigenome; (B) IDH-mutant non-codel glioma samples belonging to the group that presents intermediate shifts in the epigenome profiling; (C) IDH-mutant codel and (D) IDH-wildtype that did not change significantly in terms of their epigenome.

**Supplementary Figure 5.** Differentially methylated CpG probes between G-CIMP-low primary tumors and G-CIMP-low first recurrent tumors suggests similar patterns of epigenetic markers (n=58 hypermethylated probes and n=26 hypomethylated probes in G-CIMP-low recurrent tumors; FDR < 0.05, absolute difference in mean methylation beta value > 0.2).

**Supplementary Figure 6.** Mutational profiles of RB and AKT-mTOR pathways across a spectrum of TCGA primary gliomas. Patients are stratified based on supervised RF DNA methylation subtype, epigenomics shift profile, and risk to G-CIMP-low progression. Mutations and copy-number data were obtained from Ceccarelli et al., 2016, and visualized using cBio portal.

## SUPPLEMENTARY TABLES

**Supplementary Table 1.** Patient cohort characteristics for each patient in the cohort including clinical evolution, treatment, and molecular features.

**Supplementary Table 2.** Summary of data platform availability in each patient in the cohort. **Supplementary Table 3.** List of 712 differentially methylated CpG sites between G-CIMP-high primary tumors and their G-CIMP-low first recurrent counterparts (Figure 3A).

## ADDITIONAL RESOURCES

HOMER_Motif_Scan.zip -HOMER motif scans for the 712 differentially methylated CpG probes associated with malignant recurrence to G-CIMP-low.

Rearrangements.xlsx -A summary of DNA and RNA rearrangements organized by TCGA patient.

